# Pain distribution can be determined by classical conditioning

**DOI:** 10.1101/2024.08.31.609921

**Authors:** Jakub Nastaj, Jacek Skalski, Daria Nowak, Natalia Kruszyna, Przemysław Bąbel, Tibor Szikszay, Kerstin Luedtke, Rafał Gnat, Wacław M. Adamczyk

## Abstract

Chronic widespread pain (CWP) - as many other clinical presentations - manifests in ongoing pain without identifiable structural cause, with pain that spreads over multiple body areas. The development and maintenance of symptoms may involve learning mechanisms. Ninety-four healthy volunteers participated in this study and were randomly distributed to four groups. In the classical conditioning combined with verbal suggestion group, US-(small pain distribution) and US+ (large pain distribution) were paired with visual stimuli (CS+ and CS-) and participants were told about this association. In the verbal suggestion group, the conditioning was not performed, whereas in classical conditioning only group, learning was not combined with suggestion. In the control group, conditioning and suggestion did not take place. Ratings of perceived pain distribution (PD) were collected after each trial and ratings of pain intensity after each block of trials. During the testing phase, participants were exposed to electrocutaneous stimuli corresponding to only the small (US-) pain distribution. Results showed significant differences between CS+ and CS-pain distribution ratings across the experimental groups: conditioning + verbal suggestion (*p*<0.01), conditioning only group (*p*<0.05) and verbal suggestion only group (*p*<0.05), but not in the control group (p>0.05). Furthermore, significant differences in the perceived pain distribution were found between the control group and all experimental groups. This result supports our main hypothesis that the perceived pain distribution can be influenced by classical conditioning as well as verbal suggestion, although the effect is stronger when both are combined.

## 1. Introduction

Pain perception requires the integration of spatial and temporal aspects of sensory processing [17,43,51] and it can be experienced even in the absence of current or persistent tissue damage [46]. Many clinical presentations manifest themselves in ongoing pain without evident structural pathology. Examples include chronic non-specific pain states such as chronic widespread pain (CWP) [33], characterized by ongoing and spreading musculoskeletal diffuse pain lasting for 3 months or longer [23,33]. To develop effective treatment strategies for CWP, it is crucial to understand the mechanisms that contribute to its onset and persistence, which could involve learning mechanisms.

Acute pain is typically localized to the affected region of the body and is initiated by the activation of primary afferent neurons. This circumstance provides an optimal environment for the mechanisms underlying pain learning, particularly classical conditioning. Evidence supports the modulation of pain through learning processes such as classical conditioning [6,32,38], operant conditioning [5,37] or observational learning [49,50]. Previous studies have only examined the learning mechanisms associated with the sensory-discriminative (intensity) and affective (unpleasantness) dimensions of pain. Results showed that intensity [38] and unpleasantness of pain [45] can be amplified and perhaps even induced [15,16,27] through learning [32], sparking a heated debate of whether pain itself can be “conditioned” [20,21,28]. It is noteworthy, however, that no study has yet investigated whether the distribution of pain can be influenced by learning processes. Given the substantial published data on pain intensity [7,9,24], the authors have hypothesized that pain location could likewise be subject to classical conditioning when visual stimuli are applied as conditioned stimuli (CS). If the distribution of pain indeed originates through learning mechanisms, this can offer new perspectives accounting for CWP and spatial phenomena such as the radiation of pain [2] and spatial summation of pain (SSp) [48].

In most former studies, classical conditioning was paired with verbal suggestion to evoke a nocebo effect. Results of a meta-analysis on nocebo hyperalgesia [41] suggest that this effect is more powerful, when it arises from verbal suggestion combined with conditioning, as opposed to either verbal suggestion or conditioning alone. Moreover, generalization has been suggested as one of the mechanisms that could play a substantial role in the onset of chronic pain [39]. So far, stimulus generalization has been shown to contribute to the expansion of conditioned hyperalgesia [29,31] and analgesia [26,31] effects. However, it remains to be explored whether the spatial expansion of pain is similarly prone to stimulus generalization.

The main aim of this study was to investigate whether classical conditioning may be involved when “learning” the reported pain distribution, i.e., to address whether non-nociceptive conditioned stimuli (e.g., visual cues) can induce a pain sensation with a different (larger/smaller) reported pain distribution through classical conditioning. The secondary aim of this experiment was to investigate whether the impact of conditioning is heightened when learning is coupled with a verbal suggestion. The final aim of the experiment was to investigate whether the conditioned pain distribution extends to a perceptually similar but novel stimulus through stimulus generalization.

## 2. Methods and materials

### 2.1. Overview of the experiment

The experiment utilized a within-and between-subjects design and involved healthy volunteers assigned to one of four groups: classical conditioning only, verbal suggestion only, conditioning + verbal suggestion, or control group. Each group received the same number and type of stimuli: electrocutaneous stimuli (as unconditioned stimuli, US) inducing pain of small (US-) or large distribution (US+), which were preceded by visual (color) stimuli (as conditioned stimuli, CS). The groups differed from each other in the associations between visual and electrical stimuli. In the conditioning groups, the stimulation inducing pain of a large distribution, was consistently paired with one specific color presented to participants, whereas in the groups without conditioning, there was no association and colors were presented randomly. In the suggestion groups, participants were informed that one specific color was associated with pain distributed over a larger area. The main outcome measure in this experiment was the size of the painful area assessed via a pain distribution (PD) rating task.

### 2.2. General information

The study protocol was approved by the ethics committee at the Academy of Physical Education in Katowice (no. 4/2022) and registered at the Open Science Framework platform (https://osf.io/3qezt) using the AsPredicted.org template. The study was conducted in the certified (ISO) Laboratory of Pain Research. The experiment followed the recommendations of the Declaration of Helsinki [53]. Participants were given written and oral information of the study procedures before informed consent was obtained. Moreover, participants were informed that they can withdraw from the study at any timepoint without any reason and consequences. Each volunteer received financial remuneration for their participation. Data supporting the analyses are in reposytorium: https://osf.io/ba93f/

### 2.3. Study population and eligibility

A group of 97 healthy participants aged between 18 and 33 years (mean age 21.44 (SD 2.19) years) was recruited (between February and December 2023) **(Appendix)**. The following eligibility criteria were applied: pain free on the day of the study, healthy (self-report), no color vision deficiency, no chronic pain or experienced prolonged pain in the last 3 months, no pregnancy, no cardiovascular or neurological disease, no chronic medication use, no mental illness or any systemic disease, no electronic devices in or at the body or unremovable metal objects in the area of the non-dominant hand, as well as no skin allergies, skin lesions, tattoos or sensory abnormalities in close proximity to the non-dominant hand.

### 2.4. Sample size calculation

The required sample size was estimated based on conditioning effects observed in previous analyses of the reported pain intensity [8], which showed that at least n = 15 were needed to detect a within-group effect (pain in CS+ vs. CS-condition) of *d*_z_ = 0.92 (mean of 0.35 ± 0.38) with 90% of power and α set to 0.05 (G*Power 3.1 software) with *t* test contrast. However, given the novelty of the procedures and the need to counterbalance factors such as the color of CS (green, blue) and localization of US (proximal, distal), it was preregistered *a priori* to increase the target sample size to 24 per group. Such sample size also ensured that at least 90% of power will be maintained even with potential dropouts.

### 2.5. Outcome measures

Main outcome measure in this experiment was the length of PD expressed in centimeters – participants rated the length of the spatial dimension of pain they perceived by drawing a horizontal line on a computer screen. The longer the line, the greater the PD. The secondary outcome was the rating of pain intensity, which was collected by using a Numerical Rating Scale (NRS), ranging from “0” (no pain) to “10” (worst pain imaginable). Intensity ratings were collected to ensure that the stimulation was noxious. Further outcomes collected to characterize the study sample included: age, sex (assigned at birth), handedness, height (cm) and weight (kg). Also, fear of pain as a state was measured on a “0” (no fear at all) to “10” (highest possible fear) NRS scale and as a trait via the Fear of Pain Questionnaire III (FPQ-III) [35].

### 2.6. Apparatus, stimulation, and experimental setup

During the experiment, participants sat upright at a desk, the non-dominant hand with the electrodes attached, was placed on the desktop (with the electrodes facing the desktop). Two planar-concentric (∅ 8 mm) electrodes separated by 4cm (WASP electrodes, Brainbox Ltd., Cardiff, United Kingdom) were attached to the ulnar edge of the non-dominant hand such that they were positioned within the C8 dermatome. The dominant hand was used to perform the PD rating task (via a computer mouse) and to assess pain intensity on the NRS scale using the keyboard. Visual stimuli, PD rating task, NRS scale and written instructions were displayed on the screen (E900, BENQ, 1280×1024) placed in front of the participant (distance approximately 60 cm). Visual stimuli (colors) served as conditioned stimuli (CS) and were presented on a slide displayed on the screen for two seconds. Slides could be either blue (RGB: 19, 19, 236), green (RGB: 19, 236, 19) or cyan (RGB: 19, 236, 236). The choice of colors was driven by the results of a recent study demonstrating that these colors do not impact pain perception [52].

To induce pain, electrocutaneous stimuli were generated via a DS8R constant current stimulator (Digitimer, Welwyn, Garden City, England) with a capacity of 0 to 100mA with a maximum voltage of 400V. Each single stimulus was formed by a series of five (rectangular) pulses with 100μs duration (interval 100ms) to reduce the intensity of the current applied to the skin. Stimulation was distributed through the D188 remote electrode selector (Digitimer, Welwyn, Garden City, England) activating the required electrode (proximal OR distal – depending on counterbalanced assignment) or pair of electrodes (proximal AND distal). External control of the DS8R and D188 was ensured through a digital/analogue converter device Labjack U3-LV (LabJack Corporation, Lakewood, CO), which was controlled using “u3” Python library. Experimental procedures were fully automatic and operated by the PsychoPy (v2021.2.3) software [40].

### 2.7. Study procedures

The experiment started from the briefing and consenting procedure. The examiner verified the inclusion and exclusion criteria and prepared participants for the study procedures. The study procedure consisted of five phases: i. familiarization to electrical stimulation, ii. calibration, iii. scale training, iv. main phase of the study (manipulation depending on the group assignment), v. exit tasks.

#### 2.7.1 Familiarization

During the familiarization phase, participants received 5 electrocutaneous stimuli of different intensities in a pseudorandom order (5mA, 20mA, 15mA, 10mA and 25mA), applied using a single electrode, to familiarize participants with the sensation of electrocutaneous stimulation. This form of pre-exposure was employed to make the subsequent calibration phase and its results more valid.

#### 2.7.2 Calibration

To determine individual levels of noxious stimulus intensity for each participant, a calibration procedure was conducted. Although this procedure does not eliminate individual differences in pain perception [14], it was critical for the current experiment. Two electrodes were calibrated separately to control for the possible differences in reported pain intensity due to slightly different localizations of measurement sites. For each electrode, the method of limits was used [4] to determine the sensory detection threshold (t), pain threshold (T) and moderate pain sensation (T5) that corresponded to pain at the level of 5 on NRS scale. Electrocutaneous stimuli were delivered increasingly by 2mA increments until the participant reported the first innocuous sensation (t) and first painful sensation (T). The procedure was continued until the participant reported a moderate sensation of pain (T5) with a safety cut-off value set to 40 mA (to avoid prolonged skin irritation). Participants were excluded from further participation in the experiment if the stimulus intensity reached 40mA and participants did not report a moderately painful sensation (T5). The final intensity of stimulation (in mA) used in the main phase of the study was the same for each of the two electrodes. The final intensity(*I*_used_) was determined as the average intensity inducing moderate pain (T5) via each of the two electrodes: *I*_used_ = (T5 value for proximal electrode + T5 value for distal electrode) / 2.

#### 2.7.3. Training phase

To familiarize participants with the use of the PD rating task and to practice how to provide pain intensity ratings via the NRS, they took part in a short training phase: two noxious stimuli were applied separately, preceded by two visual stimuli to first practice the PD rating task and secondly the rating of pain intensity by using the NRS scale.

#### 2.7.4. Main phase

The main phase of the study had two components: the “experience” phase and the “testing” phase (**Figure 1A**). Participants in each group were first exposed to three blocks of 22 trials each (5 minutes break between blocks) in which CS displayed to participants were followed by a small or large pain distribution modelled here as the activation of one electrode (US-, small distribution of pain) or two electrodes (US+, large distribution of pain). Whether pain was induced via the proximal or distal electrode in the US-condition was counterbalanced across individuals.

**Figure 1.**
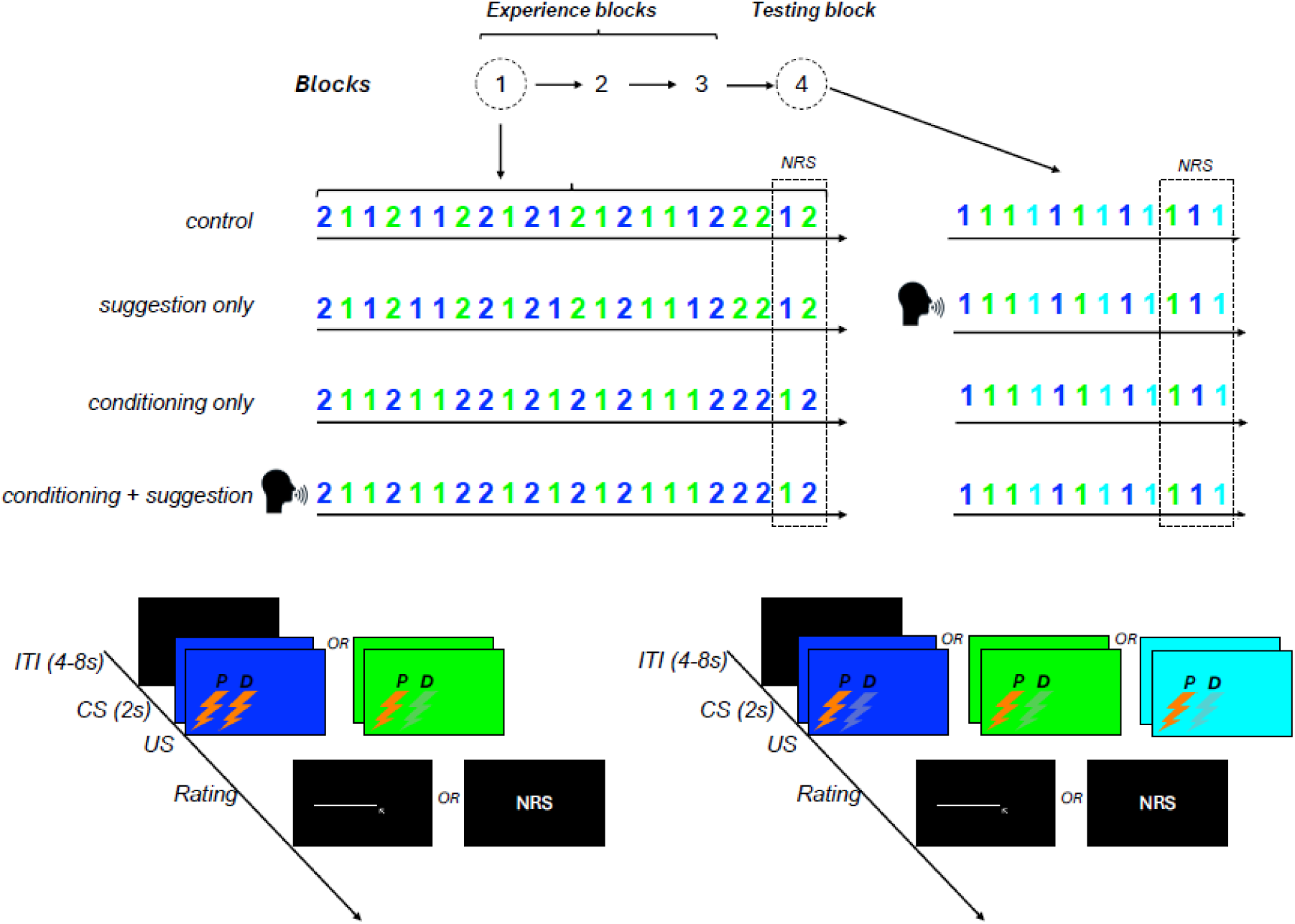
Study structure. (A) The study involved three experimental groups and one control group. Each group received the same number of nociceptive stimuli and the same number of electrodes activated for each participant (”1” or „2” indicates the number of activated electrodes). During the “experience” phase, participants underwent three blocks of trials during which noxious stimuli were delivered to them. Each block consisted of 20 trials where participants rated pain distribution using pain distribution rating task and 2 additional trials (always the last ones) where they rated pain intensity using a Numeric Rating Scale (NRS). In the “control” and “suggestion only” groups, the experience blocks featured pseudo-conditioning: the color-based conditioned stimuli [CS+ (green or blue), CS-(green or blue)] were not systematically associated with the unconditioned stimuli (US). Specifically, CS+ was not consistently paired with stimulation using two electrodes (US+), nor was CS-consistently paired with stimulation using one electrode (US-). In the classical conditioning groups, CS+ was always linked to US+ (stimulation with two electrodes), and CS-was always linked to US-(stimulation with one electrode). Of these, only one group received verbal suggestion (indicated as a pictogram) and was informed about the US-CS contingencies. During the testing block, only US-(stimulation with one electrode) was applied regardless of the CS presented. Additionally, a novel stimulus [CSGEN (cyan)], which had not been used during experience blocks was introduced to test for stimulus generalization. The “suggestion only” group received verbal suggestion (indicated as a pictogram) about the US-CS contingencies before the testing block. (B) Each given trial from experience blocks was structured with the following components: an inter-trial interval (ITI), a CS presentation period (2s) linked with US stimulation (lightning), and a rating period. During the rating period, participants either rated the pain distribution (PD) or pain intensity. To rate pain distribution participants were drawing a digital line on the screen to match the length of the pain distribution with the length of the line. Pain intensity was rated on the NRS scale displayed on the screen. (C) The trial design from the testing block was similar to the experience block, however, here always only one electrode was activated regardless of CS presentation and one new stimulus (CSGEN) was mixed with CS+ and CS-.

Depending on the group allocation, green and blue slides in this “experience” phase were consistently matched with a given US type or not. The sequence of trials was pseudorandom across each block and assignment of the colors to the US stimuli was counterbalanced across participants. For instance, green CS+ was coupled with US+ and blue CS-was coupled with US-in groups with conditioning. In other groups consistent associations were not implemented (control group, suggestion only group). Each block ended with one pair of trials in which instead of PD, pain intensity was assessed via NRS.

In the testing phase, participants were exposed to 12 trials in which always one electrode was activated (US-) regardless of the color of the CS. Apart from CS+ and CS-, which were always displayed in the “experience” phase, a new visual stimulus (CS_GEN_), i.e. cyan color, was also presented to investigate the possibility of stimulus generalization of conditioned PD. The order of CS presentations was pseudorandom. The pseudorandomization procedure was chosen to ensure that participants had equal exposure to each type of conditioned and unconditioned stimuli.

##### 2.7.4.1. Experimental manipulation

In the control group, there was no association between visual stimuli (CS+ and CS-) and pain distribution (US+ and US-) during the three blocks of trials. Activation of one (US-) or two electrodes (US+) was in pseudorandom order and the order of CS+ and CS-stimuli was also pseudorandom. In the final testing block, participants were always stimulated with one electrode (US-). No suggestion about an association between visual stimuli and distribution of pain was given to participants.

In the verbal suggestion only group, the procedure was the same as in the control group, but after the first three blocks, verbal suggestion was given before the testing phase to investigate the effect of suggestion. Associations between visual stimuli (CS+, CS-) and unconditioned stimuli (US-, US+) were suggested to participants whereas in fact, always one electrode was activated (US-). The suggestion was: *“During this phase of the experiment, after seeing the color (blue/green), you will feel pain in a smaller area, and after seeing the other color (blue/green) you will feel pain in a larger area”.* Verbal suggestion was given to the participants in written (displayed on the screen) and oral form, simultaneously. The assignment of the colors (blue, green) as CS+ and CS-was counterbalanced across the group.

In this classical conditioning + verbal suggestion group, the first three blocks consisted of trials in which stimuli leading to a small distribution of pain (US-), were paired with CS-stimuli and stimuli leading to a large distribution of pain (US+), were paired with CS+ stimuli. Verbal suggestion was given before the first of the three blocks to facilitate learning through classical conditioning and was as follows: *“Before receiving electrical stimuli, you will see a color on the monitor. After seeing the color (blue/green), you will feel pain in a smaller area, and after seeing the other color (blue/green), you will feel pain in a larger area”.* Verbal suggestion was given to the participants in written (displayed on the screen) and oral form, simultaneously. The assignment of the colors (blue, green) as CS+ and CS-was counterbalanced across the group.

In the conditioning only group, during the three blocks of trials, the small (US-) distribution of pain was paired with CS-visual stimuli and the large (US+) pain distribution was paired with CS+ visual stimuli. The assignment of the colors (blue, green) to CS+ and CS-was counterbalanced across the group. No suggestion about the association between visual stimuli and the distribution of pain was given to participants.

##### 2.7.4.2. Single trial and block design

Each trial was formed by four distinct periods which are also illustrated in **Figure 1B**: i. jittered inter-trial-interval (ITI) period (4 – 8s), ii. CS presentation on the computer screen (color slide), iii. delivery of the US, iv. rating period. Conditioned stimuli were slides of two colors, i.e. green or blue, which were associated with one of the US types, i.e. electrical stimulus applied to one electrode (US-) to model pain distribution at a small area or two electrodes (US+) to model pain of a large distribution. In the rating period (iv), participants were asked to rate the length of their pain as a proxy measure of the pain size by using the PD rating task. However, in the last two trials of each block, participants were asked to rate pain intensity by using NRS.

#### 2.7.5. Exit phase

After the main phase of the study, participants were asked to fill the exit questionnaire with five manipulation check questions. Questions were related to the study aim, meaning of the visual stimuli, contingencies between colors and reported pain.

### 2.8. Preprocessing and statistical analyses

Data saved in a raw format via PsychoPy software, were first preprocessed using MATLAB. Relevant data from the conditioning and testing phases were extracted and re-scaled to real cm values (PD ratings). All analyses were performed using R Statistical Software (v4.1.2; R Core Team 2021) [44] accepting an α-value of 0.05.

Primary analyses were performed in line with the *a priori* released study protocol (https://osf.io/3qezt). First, a mixed model ANOVA was performed on the primary outcome “PD” with “group” as the between-subject factor and “condition” (CS+, CS-, CS_GEN_) as the within-subject factor. In case of significant interaction effects, planned within-subject contrasts were performed to address specified predictions: Paired *t* test contrasts between CS+ and CS-were applied to test the outlined hypotheses: effects in groups with conditioning only, and conditioning with verbal suggestion. It was assumed that no effect will be observed in the control group and there was no specific hypothesis regarding the group with verbal suggestion only. Similar series of paired *t* test contrasts between CS-and CS_GEN_ were performed to test for the presence of stimulus generalization effects within each of the groups.

Additionally, mean within-subject main effects (differences between CS-and CS+ conditions) as well as generalization effects (differences between CS-and CS_GEN_ conditions) from experimental groups were contrasted with the effects from the control group. Lastly, for the purpose of exploratory analyses, the same set of analyses was performed on pain intensity ratings. To investigate, if evoked effects observed in the experimental groups were contaminated by potential spatial summation of pain (higher pain intensity in CS+ vs. CS-), Pearson’s correlation coefficients were calculated between PD and pain intensity. Between-group differences in descriptive statistics were explored by using one-way ANOVA with “group” as the between-subjects factor. The reliability of PD task outcomes was tested using intraclass correlation coefficients (ICC model 3,1) on the three repeated measurements for CS+, CS-, and CS_GEN_ from the testing phase of the experiment.

## 3. Results

Two participants were excluded from the study because the required intensity of reported pain was not reached during the calibration phase (5/10 on the NRS scale) within a preset intensity range (40mA). Data from one participant was not included due to technical problems during the study. One participant in the conditioning only group did not complete exit questionnaire. A total number of 94 datasets from 94 participants who completed the experiment were used to run the statistical analysis. Descriptive characteristics are shown in **Table 1**. No significant between-group differences regarding these variables were found **(Appendix).** Most of the participants in the conditioning with suggestion group and in the conditioning group noticed the association between conditioned stimuli (CS+, CS-) and unconditioned stimuli (US+, US-), contrary to participants in the verbal suggestion only group and the control group. The list of manipulation check questions and their results is presented in **Table 2**. An excellent reliability of the PD task outcomes was shown by ICC_(3,1)_ values of 0.92, 0.89, and 0.86 for the CS+, CS-, and CS_GEN_, respectively.

**Table 1.** Descriptive characteristics of the study population. F - Female, M - Male, R - right, L – left, NRS - numeric rating scale, FPQ-III – Fear of pain questionnaire, SD – standard deviation, vs – verbal suggestion

| Variable | Group |  |  |  |
| --- | --- | --- | --- | --- |
|  | conditioning + vs | conditioning | vs | control |
|  | Mean (SD) | Mean (SD) | Mean (SD) | Mean (SD) |
| Age (years) | 22.13 (2.91) | 21.09 (2.19) | 20.75 (1.57) | 21.78 (1.65) |
| Fear of pain (NRS 0-10) | 2.21 (1.79) | 1.96 (1.52) | 2.46 (1.91) | 2.13 (2.12) |
| FPQ-III: Severe pain | 29.21 (6.97) | 29.48 (7.22) | 32.17 (7.04) | 30.91 (8.15) |
| FPQ-III: Medical pain | 23.04 (6.05) | 23.13 (7.55) | 23.71 (7.19) | 24.17 (9.05) |
| FPQ-III: Minor pain | 16.92 (4.91) | 16.09 (5.96) | 16.08 (3.72) | 15.83 (5.46) |
| FPQ-III total score | 69.17 (14.95) | 68.70 (15.48) | 71.96 (14.64) | 70.91 (18.82) |
| Body mass (kg) | 72.08 (12.35) | 70.04 (11.85) | 70.21 (15.01) | 72.63 (14.59) |
| Height (cm) | 173.37 (7.73) | 172.39 (9.09) | 175.67 (8.58) | 174.09 (8.29) |
| Stimulus Intensity (mA) | 19.00 (7.48) | 16.74 (5.20) | 17.54 (5.35) | 19.00 (6.45) |
|  | **N (24)** | **N (23)** | **N (24)** | **N (23)** |
| Sex | F = 12; M = 12 | F = 13; M = 10 | F = 11; M = 13 | F = 11; M = 12 |

**Table 2.**
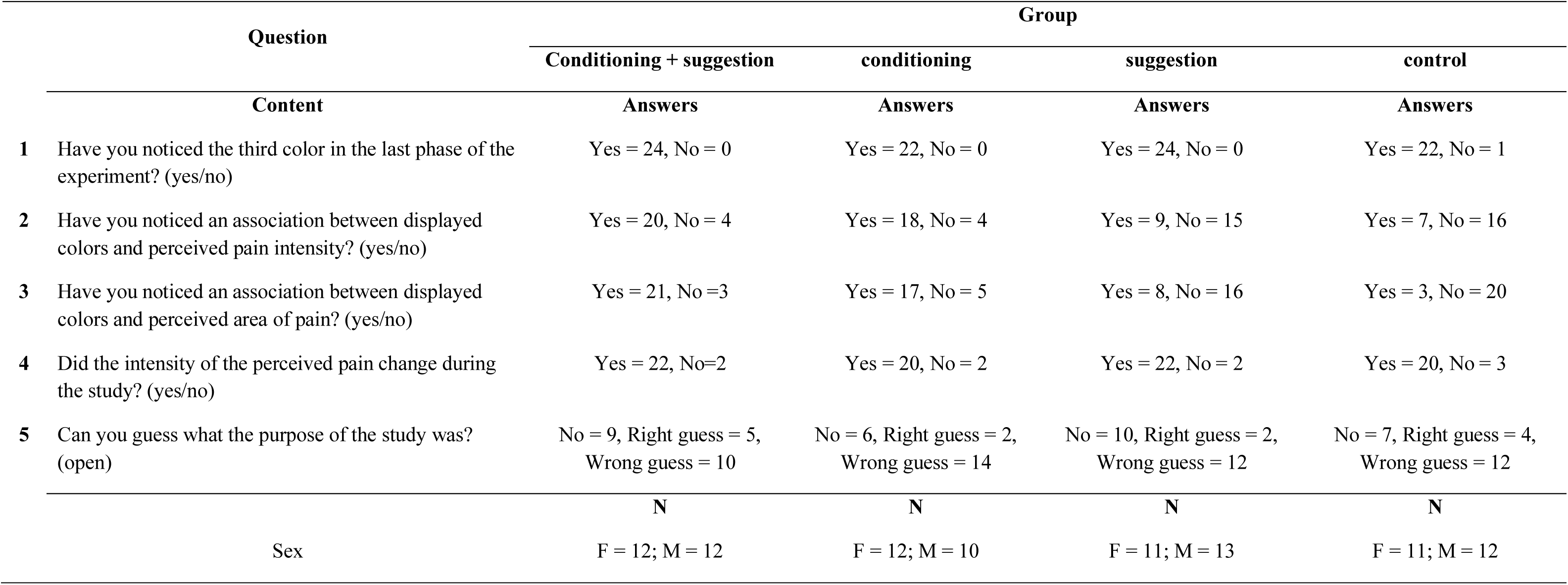
Manipulation check questions. F- Female, M -Male, Classification of answers to question 5 (open-ended): respondents could have responded with “yes” or “no”. If they chose “no” then their answer was classified as “no”. If they chose “yes” then they could answer this question in an open-ended form. If their answers included phrases suggesting the role of conditioning (learning, conditioning, predicting) or the role of suggestion (suggesting, convincing, believing) their answers were classified as “right guess,” otherwise as “wrong guess.”

|  | Question | Group |  |  |  |
| --- | --- | --- | --- | --- | --- |
|  |  | Conditioning + suggestion | conditioning | suggestion | control |
|  | Content | Answers | Answers | Answers | Answers |
| 1 | Have you noticed the third color in the last phase of the experiment? (yes/no) | Yes = 24, No = 0 | Yes = 22, No = 0 | Yes = 24, No = 0 | Yes = 22, No = 1 |
| 2 | Have you noticed an association between displayed colors and perceived pain intensity? (yes/no) | Yes = 20, No = 4 | Yes = 18, No = 4 | Yes = 9, No = 15 | Yes = 7, No = 16 |
| 3 | Have you noticed an association between displayed colors and perceived area of pain? (yes/no) | Yes = 21, No = 3 | Yes = 17, No = 5 | Yes = 8, No = 16 | Yes = 3, No = 20 |
| 4 | Did the intensity of the perceived pain change during the study? (yes/no) | Yes = 22, No = 2 | Yes = 20, No = 2 | Yes = 22, No = 2 | Yes = 20, No = 3 |
| 5 | Can you guess what the purpose of the study was? (open) | No = 9, Right guess = 5,<br>Wrong guess = 10 | No = 6, Right guess = 2,<br>Wrong guess = 14 | No = 10, Right guess = 2,<br>Wrong guess = 12 | No = 7, Right guess = 4,<br>Wrong guess = 12 |
|  |  | N | N | N | N |
|  | Sex | F = 12; M = 12 | F = 12; M = 10 | F = 11; M = 13 | F = 11; M = 12 |

### 3.1 Primary analyses: pain distribution

Mean values and standard deviations for PD and pain intensity in all groups are presented in **Table 3**. The mixed model ANOVA showed a significant effect for the factor “condition” (F_(2,180)_ = 11.35, *p* < 0.001, η_p_^²^ = 0.11) and a significant “group” × “condition” interaction was found (F_(6,180)_ = 3.54, *p* = 0.002, η_p_² = 0.11). Within-group *t* test comparisons revealed that the CS+ condition was, in general, characterized by more extended PD compared to the CS-condition (**Figure 2 and 3**). This direction was present in the conditioning + verbal suggestion group (t_(23)_ = 3.59, *p* = 0.001, *d_z_* = 0.73), conditioning only group (t_(22)_ = 2.47, *p* = 0.021, *d_z_* = 0.52), verbal suggestion only group (t_(24)_ = 2.78, *p* = 0.010, *d_z_* = 0.56) but not significant in the control group (t_(21)_ = -0.87, *p* = 0.393, *d_z_* = -0.19). Comparison between CS_GEN_ and CS-showed significantly more extended PD in CS_GEN_, however, only in the verbal suggestion only group (t_(24)_ = -2.12, *p* = 0.044, *d_z_* = -0.42). No significant differences between CS_GEN_ and CS-and stimuli were observed in the conditioning only group (t_(22)_ = -0.51, *p* = 0.612, *d_z_* = -0.11), the conditioning + verbal suggestion group (t_(23)_ = -1.93, *p* = 0.065, *d_z_* = -0.39) and the control group (t_(21)_ = 1.43, *p* = 0.166, *d_z_* = 0.31).

**Figure 2.**
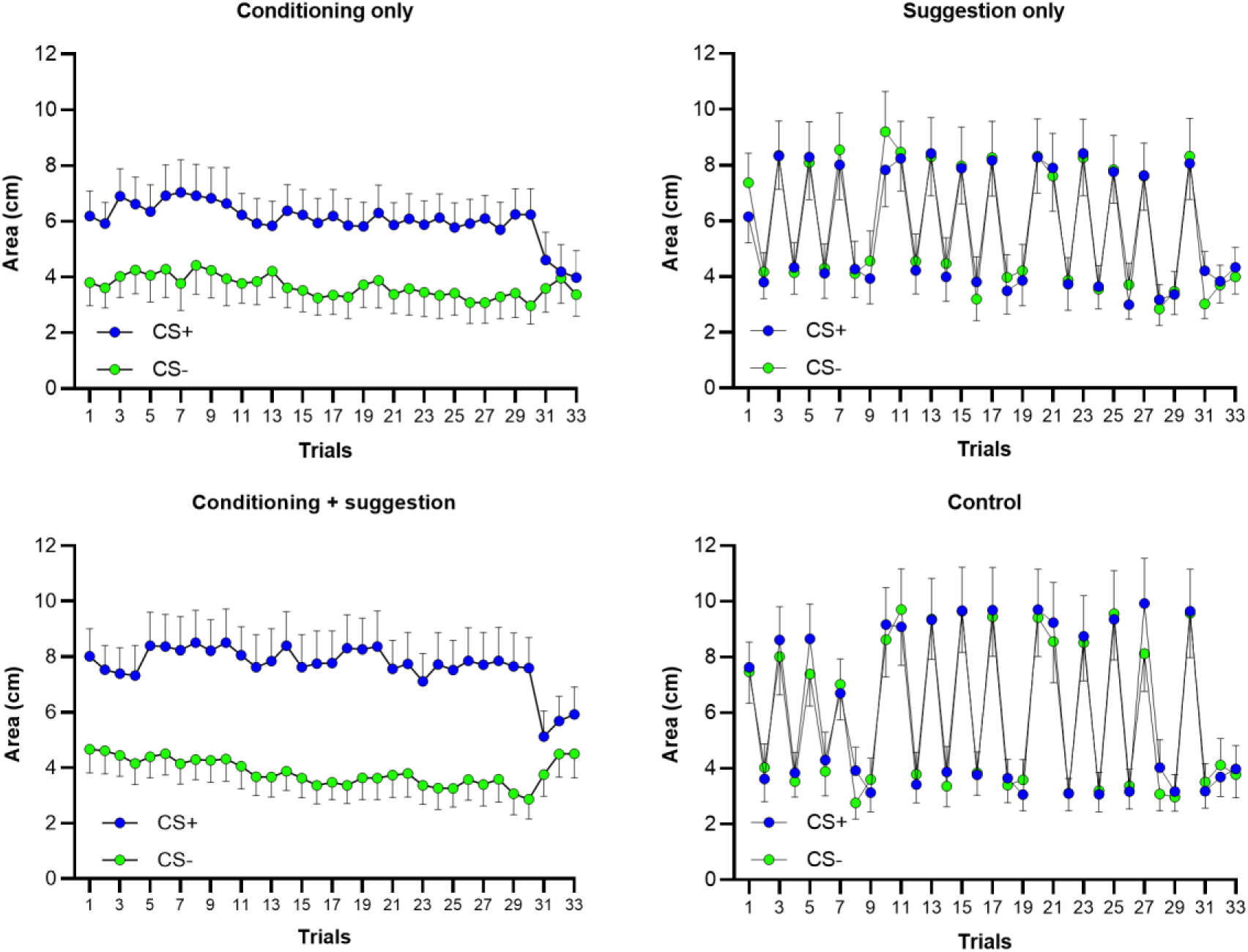
Mean ratings of reported area of pain (pain distribution) preceded by visual stimuli (CS+, CS-) in every trial. Trials: First block of experimental trials (1-10), Second block of experimental trials (11-20), Third block of experimental trials (21-30), Testing phase (30-33). CS+: Mean ratings preceded by visual stimuli associated (1-30 trials) with large pain distribution (US+) in conditioning only group and conditioning + verbal suggestion group. Mean ratings preceded by visual stimuli associated by verbal suggestion (30-33 trials) with large pain distribution (US+) in verbal suggestion only group. CS-: Mean ratings preceded by visual stimuli associated (1-30 trials) with small pain distribution (US-) in conditioning only group and conditioning + verbal suggestion group. Mean ratings preceded by visual stimuli associated by verbal suggestion (30-33 trials) with small pain distribution (US-) in verbal suggestion only group. Error bars represent the standard error of the mean.

**Figure 3.**
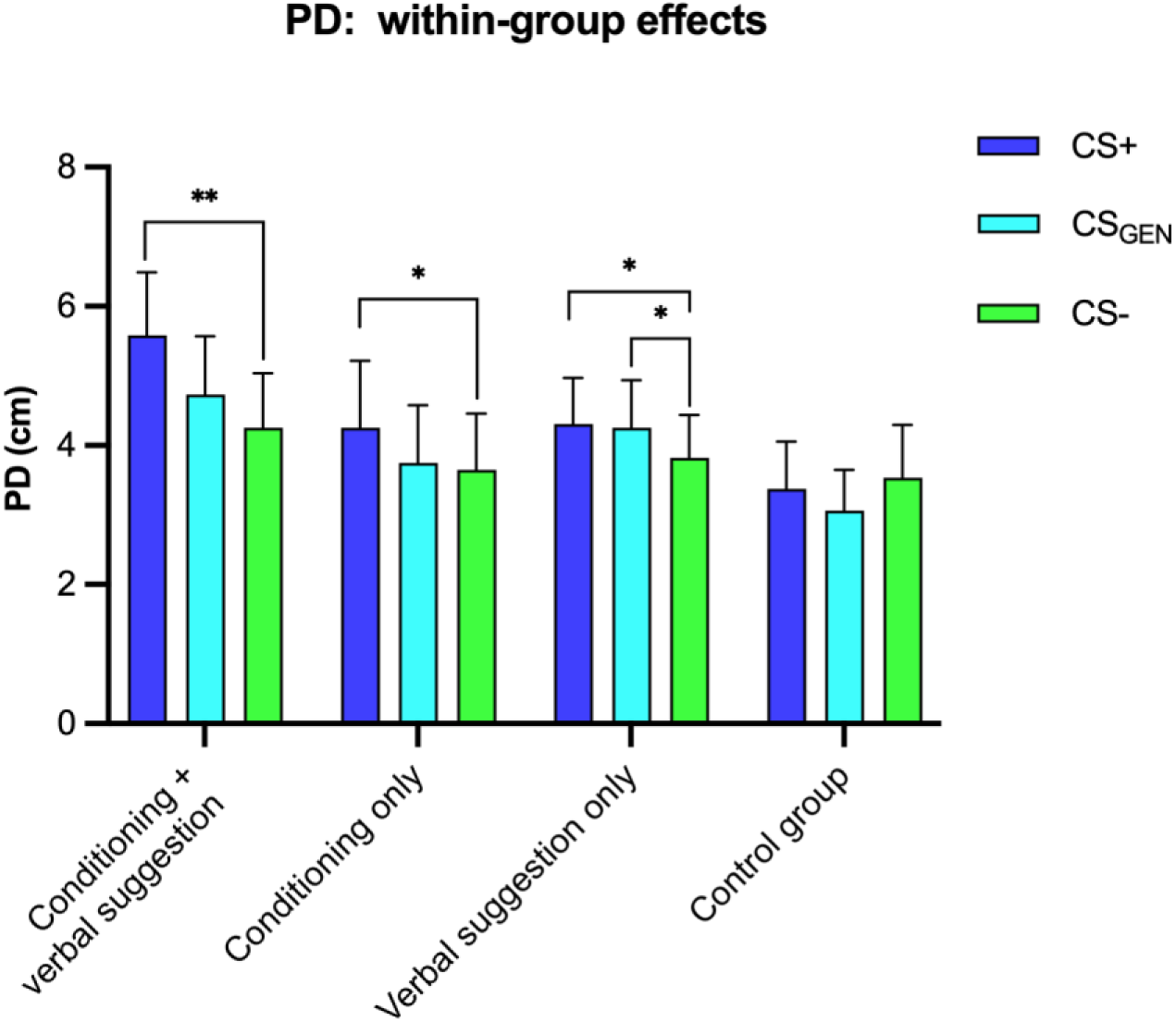
Mean ratings of reported area and intensity of pain. CS+: Mean ratings preceded by visual stimuli associated with large pain distribution (US+) in conditioning only group and conditioning + verbal suggestion group. Mean ratings preceded by visual stimuli associated by verbal suggestion with large pain distribution (US+) in verbal suggestion only group. CS-: Mean ratings preceded by visual stimuli associated with small pain distribution (US-) in conditioning only group and conditioning + verbal suggestion group. Mean ratings preceded by visual stimuli associated by verbal suggestion with small pain distribution (US-) in verbal suggestion only group. CSGEN: Mean ratings preceded by novel visual stimuli not previously associated with any pain distribution by conditioning or verbal suggestion. Error bars represent the standard error of the mean, * - p < 0.05, ** - p < 0.01.

**Table 3.** Means and standard deviations for pain distribution and pain intensity in all groups. CS+: Mean ratings preceded by visual stimuli associated with large pain distribution (US+) in conditioning only group and conditioning + verbal suggestion group. Mean ratings preceded by visual stimuli associated by verbal suggestion with large pain distribution (US+) in verbal suggestion only group. CS-: Mean ratings preceded by visual stimuli associated with small pain distribution (US-) in conditioning only group and conditioning + verbal suggestion group. Mean ratings preceded by visual stimuli associated by verbal suggestion with small pain distribution (US-) in verbal suggestion only group. CSGEN: Mean ratings preceded by novel visual stimuli not previously associated with any pain distribution by conditioning or verbal suggestion. PD-pain distribution. PI-pain intensity. SD – standard deviation. Block (1,2,3)-three blocks of trials during the experience phase. Testing – trials during testing phase.

| Outcome | Group | Block 1 |  | Block 2 |  | Block 3 |  | Testing |  |  |
| --- | --- | --- | --- | --- | --- | --- | --- | --- | --- | --- |
|  |  | CS+ | CS- | CS+ | CS- | CS+ | CS- | CS+ | CS- | CS<sub>GEN</sub> |
|  |  | Mean (SD) | Mean (SD) | Mean (SD) | Mean (SD) | Mean (SD) | Mean (SD) | Mean (SD) | Mean (SD) | Mean (SD) |
| PD | conditioning + suggestion | 8.05 (4.99) | 4.38 (3.51) | 8.00 (5.50) | 3.64 (3.24) | 7.64 (5.17) | 3.39 (3.53) | 5.58 (4.48) | 4.25 (3.83) | 4.74 (4.14) |
|  | conditioning | 6.63 (4.64) | 4.04 (4.09) | 6.07 (4.04) | 3.65 (3.62) | 6.00 (4.01) | 3.31 (3.66) | 4.26 (4.58) | 3.64 (3.91) | 3.75 (3.97) |
|  | suggestion | 5.91 (4.15) | 6.29 (4.38) | 6.04 (4.52) | 6.18 (4.63) | 5.67 (3.94) | 5.71 (4.20) | 4.32 (3.25) | 3.82 (3.00) | 4.26 (3.33) |
|  | control | 5.96 (3.59) | 5.64 (3.54) | 6.53 (4.64) | 6.56 (4.46) | 6.35 (4.58) | 6.01 (4.33) | 3.38 (3.23) | 3.54 (3.66) | 3.07 (2.82) |
| PI | Conditioning + suggestion | 4.83 (1.69) | 3.33 (2.04) | 4.92 (1.82) | 2.87 (1.62) | 4.63 (1.91) | 2.50 (1.35) | 3.88 (2.09) | 3.08 (1.74) | 3.37 (2.16) |
|  | conditioning only | 4.87 (1.60) | 2.78 (1.70) | 4.96 (1.64) | 2.48 (1.47) | 4.61 (1.64) | 2.09 (1.51) | 3.00 (1.76) | 2.48 (1.44) | 3.00 (1.65) |
|  | suggestion | 3.92 (2.04) | 3.92 (1.89) | 3.63 (1.95) | 3.92 (2.21) | 3.54 (2.28) | 3.33 (1.71) | 3.80 (1.85) | 3.40 (1.56) | 3.60 (2.12) |
|  | control group | 3.22 (1.65) | 3.57 (1.75) | 3.61 (1.80) | 3.52 (1.97) | 3.09 (1.62) | 3.13 (1.77) | 2.77 (1.41) | 2.55 (1.30) | 2.64 (1.50) |

#### 3.1.2 Exploratory analyses

To complement within-group contrasts conducted on the PD variable, additional (exploratory) between-group comparisons were performed which showed significantly larger effect sizes (difference between CS+ and CS-) in PD between the conditioning + verbal suggestion group and the control group (t_(44)_ = 3.51, *p* = 0.001, *d_z_* = 1.04), between the conditioning only group and the control group (t_(43)_ = 2.50, *p* = 0.016, *d_z_* = 0.75) and also between the suggestion only group and the control group (t_(45)_ = 2.57, *p* = 0.013, *d_z_* = 0.75).

The mixed model ANOVA performed on pain intensity, showed a more divergent pattern of results compared to the analysis of PD data. Specifically, a significant effect was found for the factor “condition” (F_(2,180)_ = 8.27, *p* < 0.001, η_p_^²^ = 0.08) but there was no main effect for the factor “group” (F_(3,90)_ = 1.92, *p* = 0.132, η_p_^²^ = 0.06) or the “condition” × “group” interaction (F_(6,180)_ = 0.78, *p* = 0.590, η_p_^²^ = 0.03), indicating that the difference between CS+ and CS-in pain intensity was consistent across all four groups. Due to the lack of effects for this outcome, no further exploratory tests were performed.

Correlations between evoked effects on PD (individual differences between CS+ and CS-) and differences in pain intensity were not significant in all groups: conditioning + verbal suggestion (r = 0.39, *p* = 0.061), conditioning only (r = 0.00, *p* = 0.997), verbal suggestion only (r = -0.04, *p* = 0.858) and control group (r = -0.26, *p* = 0.234).

## 4. Discussion

The results confirm that PD ratings preceded by CS+ or CS-stimuli were significantly different in each experimental group but not in the control group. Moreover, significant differences in PD between groups were found. Such results show that PD can be learned by classical conditioning and that the reported PD can be influenced by verbal suggestion. The largest effect size was observed in the conditioning with verbal suggestion group, which is in line with the hypothesis that verbal suggestion can enhance conditioning processes. Only in the verbal suggestion group, a significant effect of generalization was found **(Appendix)**. No significant differences in pain intensity, preceded by CS+ or CS-stimuli, were found within groups and no significant correlations were observed between pain distribution and pain intensity ratings.

### 4.1 Mechanisms of pain distribution

Pain not limited to a given body area, is a common feature of CWP. This debilitating condition is often linked with fatigue, sleep disturbances and cognitive impairment [12] and it has been proposed that peripheral as well as central mechanisms can be attributed to the widespread distribution of reported pain [11,18]. For instance, research suggests central sensitization [12,36], muscular dysfunction [10,22], bifurcation of nociceptive afferents from two different tissues [47] or deficits in endogenous pain-modulation [25,30] as some of the mechanisms explaining CWP. However, experiments on humans and animals also pointed out that experimentally induced (e.g. heat) pain is not precisely represented within the neuroaxis. Spinal cord imaging on rats demonstrated that relatively strong noxious input leads to a widespread rostro-caudal and ipsilateral-contralateral spread of neural activity within the spinal laminae that includes nociceptive-specific neurons and wide-dynamic range (WDR) neurons [13]. Interestingly, this activity spread can manifest as the radiation of pain induced by noxious heat in humans. For instance, in an experiment with two thermal probes attached to the skin and separated by 10 cm, pain was often reported under both probes even though one probe was set to deliver a neutral temperature of 35°C [42]. The most recent report with stimulation of a 2.56cm^2^ area of the skin showed that 49°C can produce radiation of pain that is, on average, 12 times greater than the size of the stimulation [2]. Thus, these psychophysical findings suggest that the human nervous system has the capacity to construct (and even interpolate) complex patterns of pain distribution based on varied sources of available nociceptive information, including population coding and neuronal recruitment [2,13,42].

It seems that the potential of the human neuroaxis to form and maintain these complex patterns of pain distribution, can be shaped by repeated interactions with the internal as well as external environments, specific to a given individual. For instance, ongoing nociception from various foci may interact with each other or with neutral environmental signals. However, this hypothetical framework for understanding the formation of pain distribution has largely been overlooked in the literature. The present study tests, if classical conditioning can contribute to the acquisition and maintenance of pain distributed at a larger area. In fact, some former translational studies have provided hints that learning can shape pain distribution. Indeed, in the study by Doménech-García et al. [19], participants who reported persistent pain in the shoulder in the past, experienced a larger area of referred pain that was triggered by a remote stimulus applied to a previously non-injured area, a pattern not observed in the healthy controls. These results may provide indirect evidence for the involvement of learning mechanisms, as additional pain stimuli could serve as conditioned stimuli eliciting a learned pain response, consistent with the findings of the present study. The engagement of learning mechanisms could account for the differences in pain distribution areas, elicited by noxious stimuli of identical intensity and applied to the same region in healthy participants.

### 4.2 Classical conditioning

The main finding of the current experiment is a demonstration that enlarged pain distribution can persist as a result of classical conditioning. Results from the primary analysis indicate a significant difference in reported PD, both between and within groups. Results from the conditioning + verbal suggestion and conditioning only group show that participants reported a wider PD in the testing phase when noxious stimuli delivered by only one electrode, were preceded by stimuli previously associated with a large PD (CS+), i.e. delivered by two electrodes, as compared to noxious stimuli preceded by stimuli previously associated with a small PD (CS-), i.e. delivered by one electrode. First, these results indicate that pain distribution elicited by electrocutaneous noxious stimulation was successfully modulated by classical conditioning. Secondly, the size of the stimulated area in the large PD (US+) was twice as large (two electrodes were active) as that of the small PD (US-) (one electrode was active) with a 2:1 proportion, indicating that a conditioned stimulus (CS+), followed by a small unconditioned stimulus (US-), can elicit pain in a larger area than the unconditioned stimuli itself. However, given the implemented method of noxious stimulation by using electrodes, the stimulated body area during the experiment was not as extensive as in many of the possible injuries from which many patients could suffer. Thirdly, differences in reported PD between CS+ and CS-were higher in the conditioning + verbal suggestion group as compared to the conditioning only group indicating that suggestion could enhance the conditioned response, here pain distribution. Such results are in line with previous studies on pain conditioning, showing a modulation of pain intensity through suggestion [41]. However, differences in the reported PD between CS+ and CS-were higher in the conditioning only group than in the verbal suggestion only group, suggesting that the learning effect could be more robust in the modulation of PD than suggestion alone, indicating the need for the first-hand experience to observe the conditioning effect.

Previous findings show that pain intensity can be modulated through classical conditioning [6,32,38]. However, no significant difference in reported pain intensity between the four groups was found in the current study. The lack of a significant difference in reported pain intensity between the groups could have been caused by the fact that authors did not manipulate pain intensity. The difference in pain intensity between a large pain distribution (US+) and a small pain distribution (US-) was the same for all groups. Furthermore, the stimuli intensity was calibrated individually. This indicates that the effect of spatial summation of pain [34] was successfully controlled and did not contaminate the “experience” phase. Moreover, a lack of any significant correlation between pain intensity and pain distribution suggests that pure pain distribution was successfully manipulated and that the results are not biased by pain intensity.

### 4.3 Limitations and future directions

This study has several main limitations. First, it only used electrocutaneous stimuli, leaving the question whether other PD modalities can be modulated by learning processes. If PD can be modulated by learning mechanisms using other types of stimuli as extensively as the intensity of pain [32], future research could significantly advance our understanding of pain “chronification”. Second, the design with two separate electrodes limits information on the extent of the conditioned responses. Future research should also explore the magnitude of the difference between US+ and US-required to condition pain sensation over a larger area, as well as the overall extent of this effect. Another unavoidable limitation of the manipulation of painful areas in this context is spatial summation of pain (SSp) [1,3], which is more intense pain with a greater area of stimulation. SSp was reduced in the current study, through the separation between electrodes < 10 cm [43], yet this did not entirely rule out the possibility of spatial summation being (to some degree) learned as a side effect. Future studies can address this through stimulus intensity adjustments. Additionally, in the control and verbal suggestion groups, the lack of a link between visual cues and pain distribution may have produced effects not seen in other groups, due to the unpredictability of stimuli. Unfortunately, manipulation check questions that were used in this study did not allow to measure such effects. Thus, it might be important to include more forms of control in future experiments, e.g. not exposing participants to a conditioning phase. Moreover, manipulation check questions should be included to measure the possible impact of stimulus predictability.

### 4.4 Conclusion

The current results confirm that pain distribution can be learned through classical conditioning and influenced by suggestion. Clinically, these findings contribute to a better understanding of the spatial features of pain, especially when patients experience pain across a more widespread area than would be expected based on the actual site of nociceptive foci. Such understanding is crucial for the development of new pain treatments. Moreover, the role of verbal suggestion in the clinical environment should not be neglected, as it can not only enhance conditioning effects but also contribute to enlarged pain distribution reports as a standalone mechanism.

## Data availability

The dataset supporting this manuscript will be made publicly available after acceptance. Funding: Polish National Science Center (2020/37/B/HS6/04196).

## Acknowledgments

The authors declare no conflict of interest. The study was funded by the National Science Centre in Poland under the grant no. 2020/37/B/HS6/04196.

