## Appendix (supplementary materials) for "Pain distribution can be determined by classical conditioning"

(supplementary materials)

### S1. Descriptive characteristics of the study sample

| Variable | Mean (SD) |
|---|---|
| Age (years) | 21.44 (2.19) |
| Fear of pain (NRS, 0-10) | 2.19 (1.83) |
| FPQ-III score | 70.10 (15.81) |
| Body mass (kg) | 71.24 (13.35) |
| Height (cm) | 173.89 (8.38) |
| Stimulus intensity (mA) | 18.07 (6.17) |
| | **N** |
| Sex | F = 47; M = 47 |
| Handedness | R = 94.7% ; L = 5.3% |

F- Female, M -Male, R - right, L – left, NRS - numeric rating scale, FPQ-III – Fear of pain questionnaire, SD – standard deviation

### S2. Between-group differences regarding descriptive statistics

| Variable | One way ANOVA | *P* value | Partial eta squared |
|---|---|---|---|
| Age (years) | $F_{(1, 92)} = 0.50$ | 0.482 | 0.005 |
| Fear of pain (NRS 0-10) | $F_{(1, 92)} = 0.05$ | 0.824 | 0.001 |
| FPQ-III score | $F_{(1, 92)} = 0.26$ | 0.608 | 0.003 |
| Body mass (kg) | $F_{(1, 92)} = 0.07$ | 0.791 | 0.001 |
| Height (cm) | $F_{(1, 92)} = 0.65$ | 0.421 | 0.007 |
| Stimulus Intensity (mA) | $F_{(1, 92)} = 0.03$ | 0.861 | 0.000 |

NRS - numeric rating scale, FPQ-III – Fear of pain questionnaire

### S3. Flow-chart of recruitment

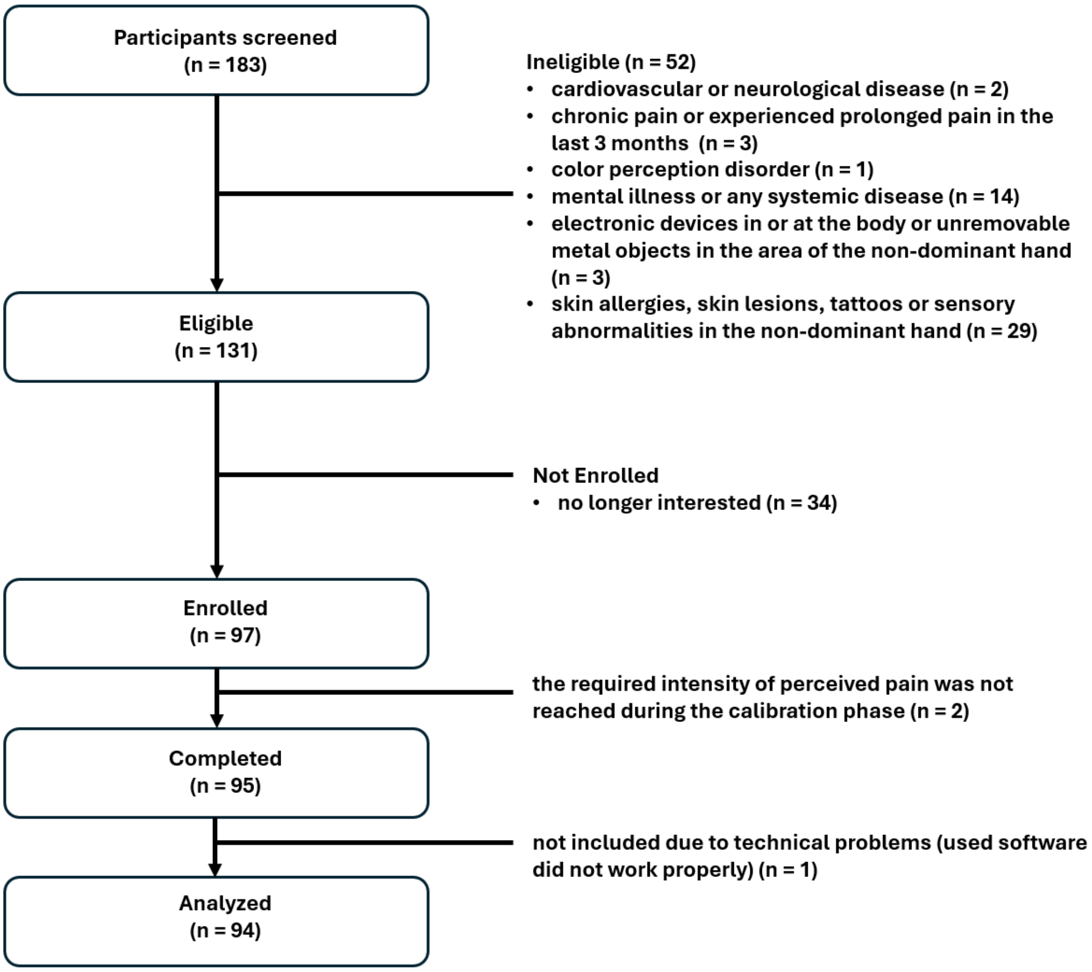

### S4. Between-group main effects (left) and generalization (right) effects

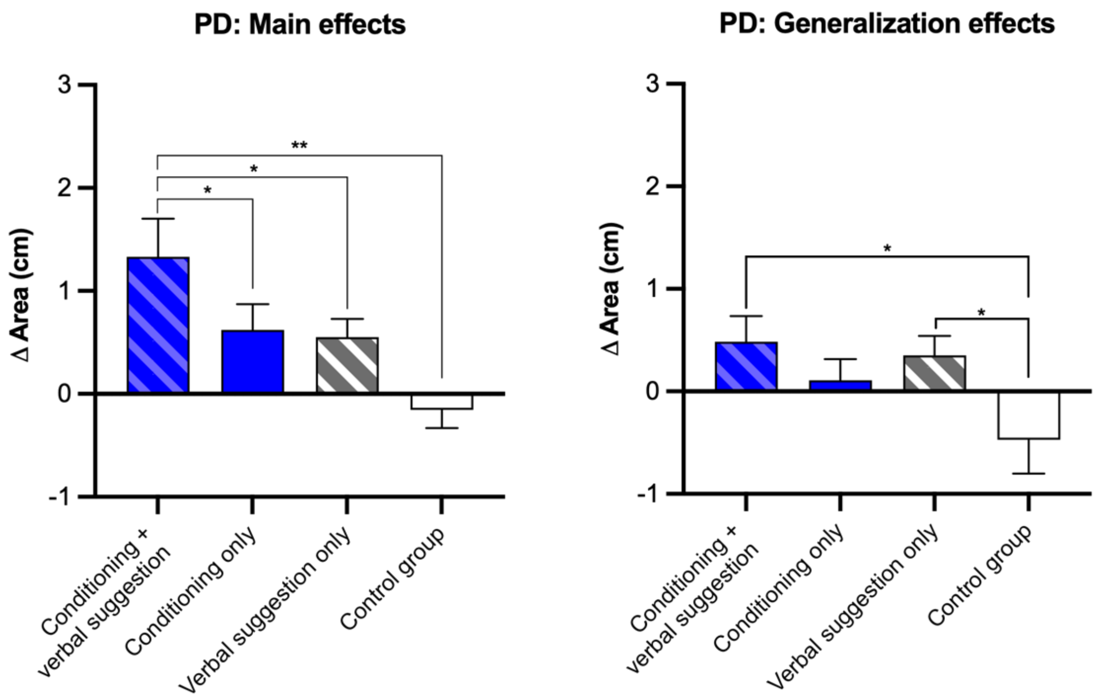


**Legend:** CS+ minus CS-: Conditioning and suggestion effect in conditioning + verbal suggestion group, Conditioning effect in conditioning only group, Verbal suggestion effect in verbal suggestion only group. $CS_{GEN}$ minus CS-: Generalization effect of mixed conditioning and suggestion in conditioning + verbal suggestion group, generalization effect of conditioning in in conditioning only group, generalization effect of suggestion in suggestion only group. PD, pain distribution. Error bars represent the standard error of the mean, * - $p < 0.05$, ** - $p < 0.01$

### S5. Discussion of generalization effects

Generalization and discrimination of stimuli and reaction are essential parts of learning through conditioning [6]. Current results do not provide evidence for a significant generalization effect of the conditioned pain distribution, only in verbal suggestion group a significant effect was observed. Therefore, a lack of generalization of the learning of pain distribution seems implausible. However, most previous studies focused on the generalization of a pain related fear reaction [2,7,8] and not on the generalization of pain intensity itself [3–5]. Thus, more evidence is needed to support the hypothesis that both pain distribution and pain intensity can be generalized as a conditioned reaction. In this experiment, the cyan color (RGB (19, 236, 236)) was chosen as the generalization stimulus because the color is embedded "in-between" blue (RGB (19, 19, 236)) and green (RGB (19, 236, 19)). Generalization depends strongly on the physical properties of similarity of presented novel stimuli to conditioned stimuli, such as the intensity of stimulation (the total activation of receptors in the sense organs), and how a given amount of stimulation is distributed among receptors [1]. As novel stimuli used in the current study were evenly "in-between" blue and green it could lack crucial physical similarities to conditioned stimuli (CS+) to elicit a generalized response as novel stimuli was evenly similar to CS+ and CS-.
